# A Framework for Evaluating Myocardial Stiffness Using 3D-Printed Hearts

**DOI:** 10.1101/2021.02.21.432169

**Authors:** Fikunwa Kolawole, Mathias Peirlinck, Tyler E. Cork, Vicky Y. Wang, Seraina A. Dual, Marc E. Levenston, Ellen Kuhl, Daniel B. Ennis

## Abstract

MRI-driven computational modeling is increasingly used to simulate *in vivo* cardiac mechanical behavior and estimate subject-specific myocardial stiffness. However, *in vivo* validation of these estimates is exceedingly difficult due to the lack of a known ground-truth *in vivo* myocardial stiffness. We have developed 3D-printed heart phantoms of known myocardium-mimicking stiffness and MRI relaxation properties and incorporated the heart phantoms within a highly controlled MRI-compatible setup to simulate *in vivo* diastolic filling. The setup enables the acquisition of experimental data needed to evaluate myocardial stiffness using computational constitutive modeling: phantom geometry, loading pressures, boundary conditions, and filling strains. The pressure-volume relationship obtained from the phantom setup was used to calibrate an *in silico* model of the heart phantom undergoing simulated diastolic filling. The model estimated stiffness was compared with ground-truth stiffness obtained from uniaxial tensile testing. Ultimately, the setup is designed to enable extensive validation of MRI and FEM-based myocardial stiffness estimation frameworks.

## 1 Introduction

Heart failure (HF) is a condition in which the heart is unable to meet the metabolic demands of the body. The US public health burden of heart failure is expected to grow significantly in the next decade with the prevalence projected to reach 8 million by 2030 [1]. As HF often occurs because of deteriorating cardiac function due to persistent remodeling, pathophysiological cardiac remodeling has been identified as a therapeutic target in HF [2]. Thus, understanding the various mechanisms and manifestations of remodeling is fundamental for formulating appropriate clinical intervention. A significant consequence of cardiac remodeling is changes to the passive (diastolic) stiffness of ventricular myocardium. Measuring *in vivo* passive myocardial stiffness requires a comprehensive evaluation of the mechanical behavior (stress-strain) in diastole.

Passive myocardial stiffness is commonly inferred from the left ventricle (LV) end diastolic pressure-volume relationship (EDPVR) [3]. However, EDPVR only provides a global estimation of the LV chamber stiffness. It is therefore inappropriate to infer intrinsic myocardial mechanical behavior from EDPVR. Cardiac Magnetic Resonance Elastography (MRE) has been used for direct measurement of myocardial shear stiffness [4], but its implementation requires assumptions that make solutions challenging for *in vivo* cardiac physiology.

Computational modeling using MRI data and Finite Element Modeling (FEM) enables the estimation of *in vivo* myocardial passive stiffness [5]. Provided ventricular geometry, microstructural organization, boundary conditions, kinematics and volumes from MRI, and endocardial filling pressures from catheterization, continuum balance laws and optimization techniques can be leveraged in FEM to inversely obtain the parameters of the constitutive model governing the material’s mechanical behavior. Currently, however, validation of MRI and FEM-based myocardial stiffness estimation frameworks is exceedingly difficult *in vivo* as ground-truth *in vivo* myocardial stiffness remains elusive.

We have developed a 3D-printed heart phantom with a myocardium-mimicking material of known stiffness and MRI properties. The phantom was incorporated within an MRI compatible *in vitro* diastolic filling setup. We estimated stiffness of the heart phantom using an MRI-derived computational model and compared with ground-truth measures obtained from uniaxial tensile testing.

## 2 Methods

### 2.1 Phantom Development and Material Characterization

High-resolution T1-weighted images from a healthy *ex vivo* porcine heart (restored to *in vivo* mid-diastasis geometry) [6] were used to generate a 3D geometric heart model. The epicardial surface of the 3D geometric heart model was used to create a negative epicardial mold. The ventricular blood pool segmentations were used to create LV and right ventricle (RV) blood pool casts. The epicardial mold and blood pool casts were converted into stereolithography files and 3D printed (Ultimaker 3 Extended) using tough polylactic acid and water-soluble polyvinyl acid, respectively. The heart phantom was cast using a silicone elastomer blend containing mass ratio of Sylgard 184:527 (Dow Corning) of 1:4 (Figure 1). The suitability of different Sylgard blends was assessed, and the tissue-mimicking material was chosen based on mechanical and MR relaxation properties. Heart phantoms were cast by curing the Sylgard blend in the 3D printed mold for 48 hours at room temperature. Ventricular basal ports and an apical anchor were added to the heart phantom to facilitate loading and motion stabilization.

**Fig. 1.**
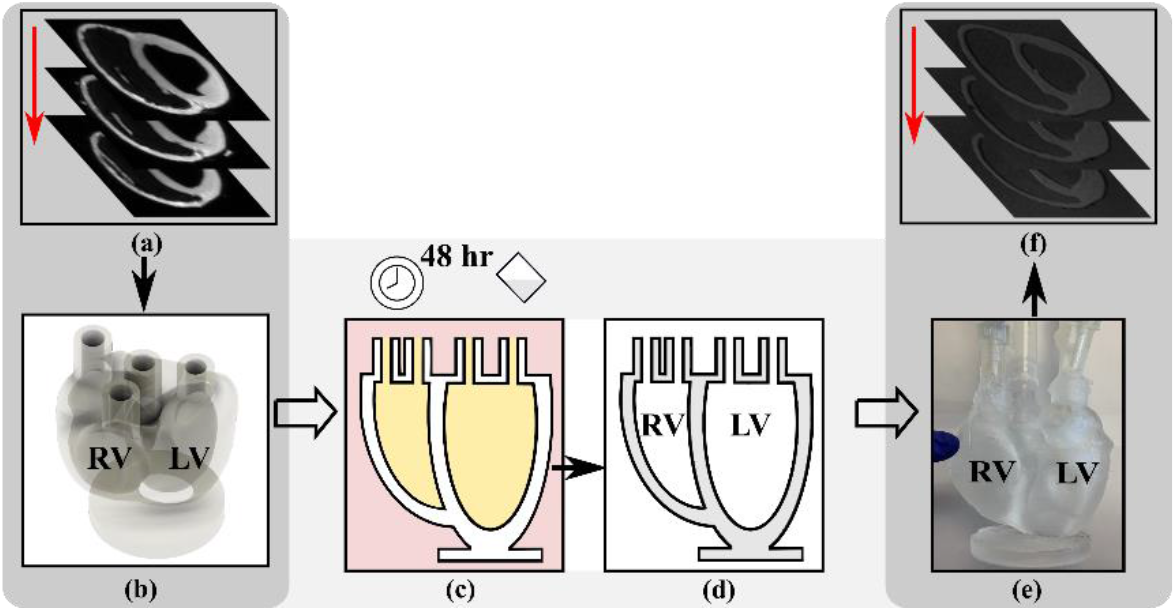
Phantom Development. (a) *ex vivo* porcine T1-weighted images segmented to develop a geometric model. (b) model fitted with ports and apical anchor. (c) Sylgard blend cured in mold for 48 hours. (d) epicardial mold (pink) removed and ventricular blood pool casts (yellow) dissolved in water. (e) subject-specific heart phantom with ports. (f) phantom 3D-SPGR images.

Uniaxial tensile testing samples of the Sylgard blend were produced in parallel with the phantom development. Samples were punched out (ASTM cutting die A) from cured sheets (thickness 3.13 ± 0.11mm) and three samples were mechanically tested according to the ASTM D412 standard using an Instron 5848 Microtester (100N load cell). Samples were mounted by first clamping the specimen to the upper grip, zeroing the load cell, then clamping the specimen to the lower grip while taking care to avoid load application. The extensometer was then clamped to the samples with the gauge length set at 50.8mm. The test was performed at ambient conditions with a strain rate of 500 mm/min. The Elastic Modulus was evaluated from a linear regression of the stress-stretch relation at stretches from 1.0 to 1.2. MRI relaxation properties were measured using T1-mapping (MOLLI 5-3-3, spatial resolution 1.00×1.00×5.0mm^3^) and T2-mapping (T2-prep FLASH; flip angle 12°; spatial resolution, 1.00×1.00×5.0mm^3^).

### 2.2 *In vitro* Diastolic Filling Setup

The final heart phantom was embedded a flow loop (Figure 2) controlled by an MRI-compatible linear motion stage (MR-1A-XRV2, Vital Biomedical Technologies). To simulate *in vivo* diastolic LV filling, the loop was designed to deliver a cardiac-like late-diastolic filling cycle in the LV (sinusoidal flow: 13 mL/cycle mean, 1s period).

**Fig. 2.**
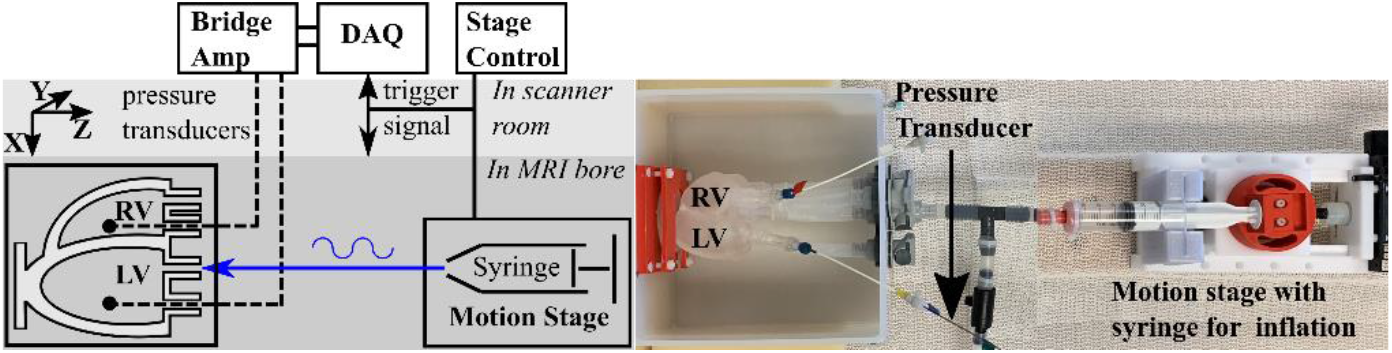
Experimental Setup. Schematic (left) and picture (right)

**Fig. 3.**
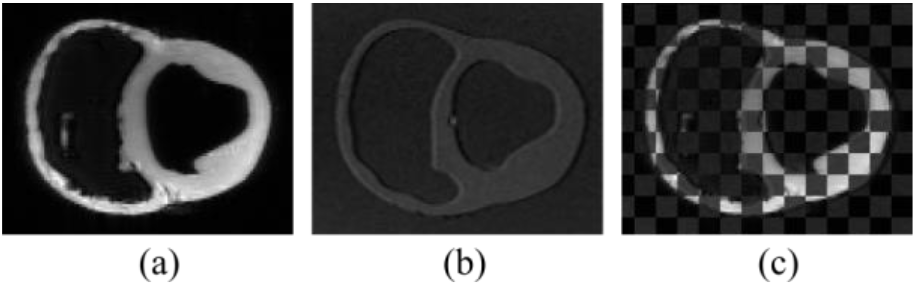
Short-Axis slice. (a) T1-weighted *ex vivo* porcine subject (b) 3D SPGR *in vitro* heart phantom (c) qualitative checkerboard of *ex vivo* and *in vitro*

Ventricular pressure was acquired continuously in PowerLab (ADInstruments) using MRI-compatible pressure transducers (Micro-Tip SPR 350S, Millar). LV filling volume was recorded from the motion stage and synchronized with pressure measurements. The RV was kept at a constant volume and low pressure to mimic the relatively low pressures and pressure variations in the RV during diastasis, compared to the LV. Each ventricle was connected in a closed loop and the fluid volume in each loop was fixed. The motion stage was used to deliver the sinusoidal flow through a fluid filled syringe within the LV loop. The phantom was fixed at the apex and basal ports.

### 2.3 Image Acquisition and Processing

All *in vitro* images were collected on a 3T (Skyra, Siemens) using a 32-channel chest and spine coil. In a static equilibrium phase, a 3D SPoiled Gradient Recalled echo (3D SPGR; TE/TR=2.17/5.5ms; FA=20°; isotropic 1.00mm^3^; Static equilibrium phase) was performed over the entire volume of the phantom.

The 3D phantom geometric fidelity was evaluated using *in vitro* and *ex vivo* images semi-automatically segmented to extract binary masks of the myocardium, LV blood pool, and RV blood pool using Otsu’s method and manual clean up (MITK Workbench). *In vitro* myocardium masks were registered to *ex vivo* myocardium using an automated 3D rigid regular-step gradient descent algorithm (Matlab, Mathworks). This transformation was applied to all *in vitro* binary masks. Volumetric accuracy and dice similarity coefficient (DSC) of binary masks were assessed by comparing the volume of the myocardium, LV blood pool, and RV blood pool from the *in vitro* images to *ex vivo* images.

### 2.4 *In silico* modeling and stiffness quantification

Starting from the 3D geometrical model (Section 2.1), a volumetric quadratic tetrahedral mesh with an average edge size of 1.5mm was constructed using the 3-Matic meshing software suite (Materialise). For computational efficiency, the constrained apical anchor was removed from the mesh. The resulting mesh consisted of 118,337 nodes and 61,272 tet10 elements, summing up to 355,011 degrees of freedom to be solved in the finite element analysis software suite Abaqus (Dassault Systemes, Simulia Corp). Previous studies on mechanical characterization of Sylgard 184 and 527 indicate that the materials are nearly incompressible, or weakly compressible (ν=0.495) [7]. Thus, we assumed near incompressibility in the Sylgard blend. Given that we are working in a lower strain regime (strains under 40%), the Sylgard silicone elastomer blend was assumed to be linearly elastic [8] (consistent with our benchtop data). In accordance with the experimental setup, the computational heart phantom was kinematically constrained by encastring the apical bottom surface and the top surfaces of the ventricular basal ports. The port openings were virtually closed off to create enclosed fluid cavities, allowing an efficient computation of the temporal pressure and volume evolutions in the left and right ventricular ‘blood’ pools. Using a volume-driven boundary condition on the left and right ventricular fluid cavities, the deformation of the phantom following the sinusoidal inflow and outflow was virtually simulated.

The resulting pressure field was used to compute the elastic stiffness of the 3D casted Sylgard blend. More specifically, we found the Young’s Modulus (E) using Abaqus as the forward solver wrapped inside Python’s Nelder-Mead optimization algorithm [9,10]. Starting from an initial value (E=150 kPa), we computed the mismatch between the experimentally measured and simulated pressure evolution, and iteratively updated the Young’s Modulus until this error was minimized.

## 3 Results

### 3.1 Phantom Geometric Accuracy

Analyses of the 3D *in vitro* heart phantom and *ex vivo* porcine subject MR images showed that the phantom development procedure adequately reproduces the porcine subject heart geometry. The *in vitro* and *ex vivo* images were all well registered.

Table 1 reports results for quantitative assessment on geometric fidelity of the *ex vivo* porcine subject heart geometry compared to the *in vitro* heart phantom. DSC is reported for the LV blood pool, RV blood pool, and myocardium. Geometric fidelity was also assessed by comparing the LV blood pool, RV blood pool, and myocardium volumes between the phantom and the swine subject.

**Table 1.** Geometry comparison between *in vitro* (phantom) and *ex vivo* (porcine subject)

| Tissue | DSC | *In vitro* volume | *Ex vivo* volume | Volume Error |
| --- | --- | --- | --- | --- |
| LV Blood Pool | 0.92 | 45.5 mL | 43.9 mL | 3.6 % |
| RV Blood Pool | 0.95 | 67.5 mL | 68.9 mL | -2.0 % |
| Myocardium | 0.89 | 90.8 mm<sup>3</sup> | 93.0 mm<sup>3</sup> | -2.4 % |

### 3.2 Uniaxial Tensile Testing

The Cauchy stress versus the stretch ratio for a representative sample of the Sylgard blend used to cast the heart phantom is shown in Figure 4a. The elastic modulus was calculated for the three samples by fitting a linear model to the stress-stretch data for stretches between 1.0 and 1.2 and estimated to be 235 ± 6kPa (mean ± SD). T1 and T2 times were 959.5 ± 5.0ms and 313.8 ± 12.3ms, respectively.

**Fig. 4.**
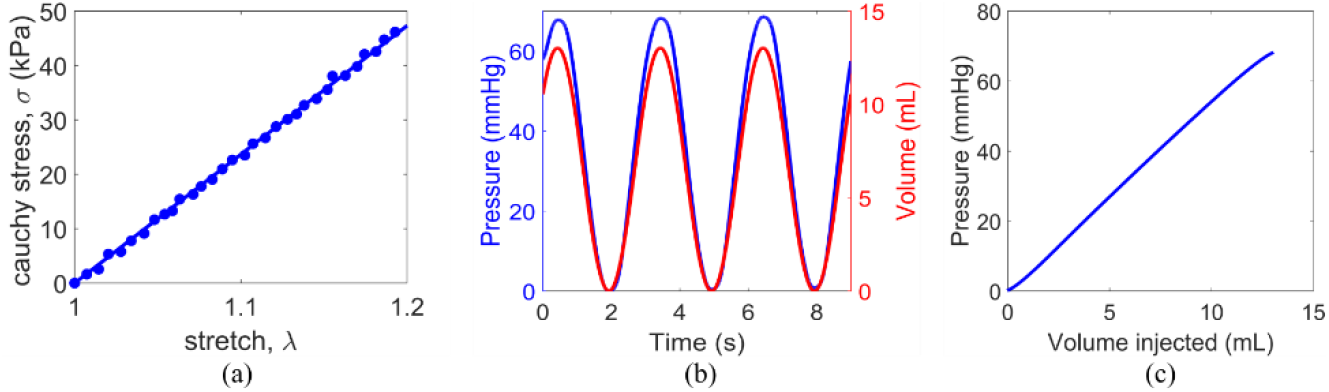
(a) Cauchy stress vs Stretch. (b) Pressure & Volume vs Time (c) *P-V* Experimental

### 3.3 Pressure-Volume

Recorded pressures were sinusoidal (mean 34 mmHg, min. 0 mmHg, max 68 mmHg) as were the volumes (peak 13mL; mean 6.5mL) (Figure 4). The LV pressure-volume (*P-V)* relation was obtained after the loading and unloading cycles were steady and repeatable. The loading *P-V* curve was used for the *in silico* stiffness calibration.

### 3.4 *In silico* stiffness calibration

Figure 5a shows the computed deformation and the spatial variation in maximum principal stretch state at the maximal loaded volume during the sinusoidal loading protocol. The maximal stretch in the phantom is located at the endocardial LV surface, reaching maximum principal stretches up to 1.2. The *in silico* stiffness calibration is shown in Figure 5b. Starting from an initial elastic modulus of 150 kPa, the elastic modulus was iteratively updated, minimizing the pressure differences between the computational (line plot) and the experimental (x-markers) results. The line plot represents the converged forward simulation from which an elastic modulus of 328 kPa was obtained.

**Fig. 5.**
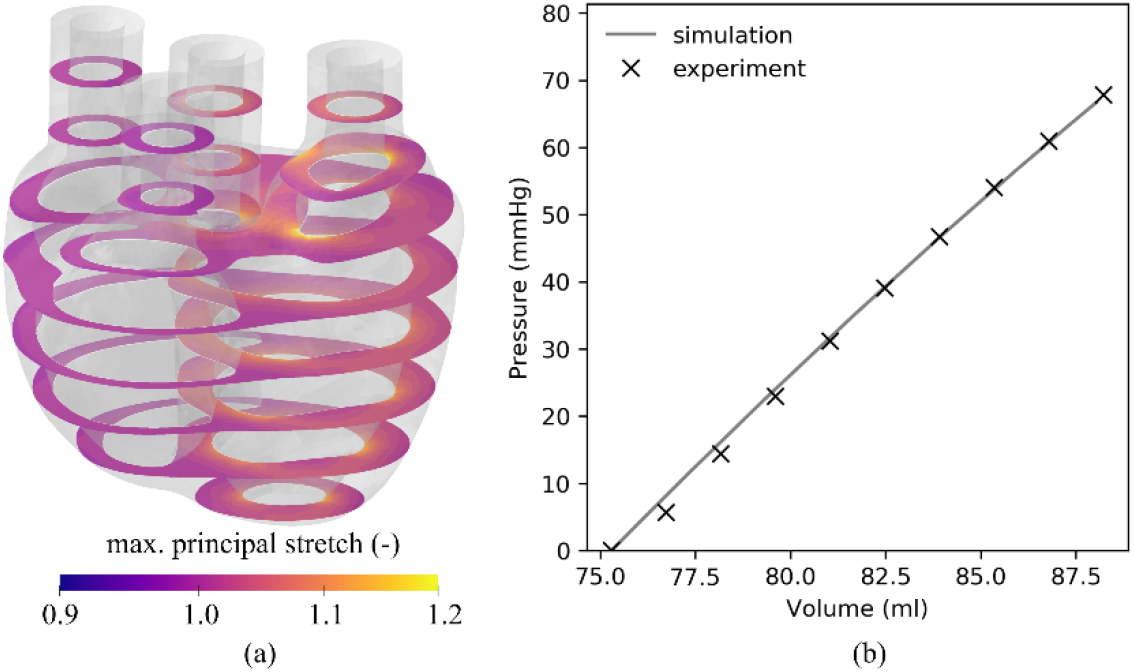
(a) Computed phantom deformation at maximal inflation. Short-axis slices depict the spatially varying stretch of the Sylgard blend. (b) Calibrated elastic modulus of 328 kPa.

## 4 Discussion

To validate our MRI and FEM-based stiffness estimation framework, we have developed subject-specific heart phantoms with passive ventricular myocardium-mimicking mechanical and MRI relaxation properties. The heart phantom adequately replicates the porcine subject geometry as shown by the reported high DSC between registered 3D SPGR *in vitro* (heart phantom) and the *ex vivo* (porcine subject, geometrically restored to *in vivo* mid-diastasis) images.

The properties of the material used to construct the phantom were comparable to myocardium, but lacked some key features. The chosen Sylgard blend is considerably stiffer than myocardium. By comparison, Sommer *et al.* in their work on *ex vivo* mechanical characterization of myocardium obtained maximum Cauchy stresses in the fiber direction of about 8 kPa at stretch of 1.1 [11]. Our material, on the other hand, exhibited Cauchy stress of about 22 kPa at the same stretch. Although Sylgard blends with more Sylgard527 are softer, this blend was chosen for its far superior workability. Additionally, the T1 relaxation time of the Sylgard blend was identified as 959.9ms at 3T compared with reported myocardium T1 times of 1158.7ms at 3T [12]. The T2 relaxation time, which is less important for our study was however, significantly higher than that of myocardium at 313.8ms compared with 45.1ms [12].

We estimated the phantom stiffness in an *in silico* model by matching the simulated compliance-pressure curve to the *in vitro* experimental *P-V* relation obtained from the heart phantom diastolic filling setup. The simulated solution converged to an elastic modulus of 328kPa compared with the phantom material stiffness ground-truth of 235 ± 6kPa obtained through tensile testing. Palchesko *et al.* characterized blends of Sylgard 184 and 527 [13]. For our blend, their experimentally derived relationship between mass composition of a blend and its elastic modulus gives a stiffness of 215 kPa, comparing favorably with results from our tensile testing (235 kPa). This, and our measurement data, suggests that our stiffness simulation is overestimating the material stiffness. The discrepancy between simulated and benchtop estimate of phantom mechanical properties could be due to inaccuracies in the *P-V* used for stiffness calibration. It is however more likely that the discrepancy is indicative of variations in mechanical properties between the heart phantom and tensile testing specimen. Although the heart phantom and tensile testing specimen were developed in parallel from the same batch, their mechanical properties may differ due to different curing conditions. The mechanical properties of Sylgard are sensitive to curing temperature and aging time between manufacture and testing [14]. This effect may be accelerated in thinner samples, which may explain variability in mechanical properties between the tensile testing samples and the heart phantom. To verify ground-truth phantom stiffness, in the future, testing strips could be cut out directly from the heart phantom after MRI and pressure data have been obtained.

Our future studies will use MRI tagging or displacement encoding to reconstruct local phantom displacements during the filling cycle. The filling pressures and kinematics will be used for *in silico* stiffness calibration, providing more constraints for stiffness estimation, thus improving the accuracy of simulated phantom stiffness.

In conclusion, we have developed an experimental setup for validating MRI and FEM-based myocardial passive stiffness in 3D-printed heart phantoms. The setup enables acquisition of MRI and pressure data necessary to estimate phantom material properties using an MRI and FEM-based constitutive modeling framework. The phantom development procedure effectively reproduces the subject geometry. The experimental setup will be refined to minimize discrepancy between the phantom mechanical properties and those of the tensile testing strips used to quantify ground-truth stiffness. The phantom stiffness can be tuned, and different subject-specific phantoms can be developed, enabling extensive quantification of the accuracy and repeatability of stiffness estimates obtained through our MRI and FEM-based stiffness estimation framework.

## Notes

### Competing Interest Statement

The authors have declared no competing interest.

